# Kūkulu: Diffusion-Based Reconstruction of Antibody CDR Loops using a Structure-Aware Joint Embedding Predictive Architecture

**DOI:** 10.64898/2026.08.03.742168

**Authors:** Seth Rabinowitz, Prbhuv Nigam, Nicholas Santolla, Colby T. Ford

## Abstract

Antibody complementarity-determining regions (CDRs), especially CDR-H3, are a dominant source of binding specificity but remain difficult to design due to coupled sequence-structure constraints and local geometric variability. Here we present *Kukulu*, a structure-aware Joint Embedding Predictive Architecture (JEPA) combined with conditional diffusion for CDR loop reconstruction in antibody-antigen complexes. Our pipeline prepares structures by chain-aware cleanup, Fv trimming, Chothia-indexed CDR identification, and *in silico* CDR masking, then trains on paired prepared/masked structures represented in an atom37 format. The model uses a context encoder over masked structures, a transformer predictor for latent CDR representations, and a diffusion head that reconstructs loop coordinates, atom presence, and residue identities under geometry-aware losses. During generation, *Kukulu* denoises only masked CDR residues while preserving framework context, then optionally rebuilds sidechains with local frame templates and performs post-generation structural relaxation. This manuscript provides a methods-focused overview of the model’s implementation details and an evaluation protocol based on structure quality and docking-oriented scoring for integration into existing antibody design workflows.

## Introduction

Antibody therapeutics have transformed treatment across oncology, immunology, and infectious disease, but prospective antibody design remains challenging because binding and developability depend on tightly coupled local sequence and three-dimensional structure in the variable domain (1, 2). Classical computational antibody design methods, including *RosettaAntibodyDesign*, established principled frameworks for structure-guided grafting and optimization but are often limited by search complexity and handcrafted energy landscapes in highly flexible loop regions (3).

Recent artificial intelligence (AI) methods have expanded the design space. Protein language models and inverse folding approaches support sequence design from structure-conditioned context (4), while diffusion-based generative models provide a route to multimodal structural sampling in proteins (5, 6). In antibodies specifically, diffusion-based approaches such as DiffAb demonstrate that targeted CDR generation can be learned with explicit structural conditioning (7). However, robust generation of antibody loops still requires preserving framework geometry, handling variable loop lengths and chain boundaries, and integrating residue identity with all-atom coordinates.

Here, we introduce *Kukulu*, a structure-aware JEPA + diffusion method for CDR loop reconstruction in existing antibody-antigen complexes. JEPA-style latent prediction supports context-conditioned representation learning without direct token-level autoregression (8), and we pair it with VICReg-style regularization to reduce representation collapse in masked CDR tokens (9). The core goal is to reconstruct physically plausible CDR loops from masked complexes while preserving framework context and producing complete atom37-compatible outputs suitable for down-stream docking and design analyses.

This manuscript emphasizes: (i) a reproducible preparation workflow from raw PDB complexes to paired training inputs, (ii) a hybrid architecture that couples JEPA latent prediction with coordinate diffusion, and (iii) a practical inference stack that supports CDR replacement, optional sidechain rebuilding, and post-generation relaxation.

## Methods

*Kukulu* models CDR loop reconstruction as a conditional generation problem: given a masked antibody-antigen complex in which CDR residues are removed, predict loop residue identities, atom presence, and all-atom coordinates for masked CDR positions. The implementation is organized into structure preparation, atom37-based tensorization, paired masked/prepared dataset assembly, JEPA + diffusion training, and post-processed inference.

### Structure Preparation and CDR Masking

The data curation process began by using the SAbDab-based dataset from our group’s *peleke-1* model training, which contains PDB IDs and each structure’s chain IDs for the heavy chain, light chain, and antigen chains (10–12). Then, raw structures were downloaded from the Protein Data Bank using *Biopython* utilities (13). Structures were cleaned chain-by-chain with *PDBFixer* (missing residue/atom handling, nonstandard residue replacement, heterogen removal, unneeded chain removal) and recombined into a single complex (14).

Fv and CDR boundaries were annotated from chain sequences with *AbNumber*/*ANARCI* using the Chothia scheme (15, 16). Structures are trimmed to the Fv region for heavy and light chains while preserving antigen chains. Prepared structures are then converted into masked inputs by removing residues within CDR-H1/2/3 and CDR-L1/2/3 bounds. The result is a paired dataset: “*_prepared.pdb” targets and “*_masked.pdb” contexts.

### Atom37 Representation and Pair Alignment

Each residue is represented in a fixed 37-atom frame (AlphaFold/OpenFold-style layout) with coordinate tensor shape *L* × 37 × 3 and atom-presence mask shape *L* × 37, where *L* is residue length. Residues are keyed by chain ID, residue number, and insertion code to robustly align prepared and masked complexes. Missing residues in masked structures define a binary CDR token mask used throughout training and inference.

Batch collation supports variable sequence lengths through right-padding and a residue mask. Additional supervision tensors include per-residue amino acid targets and chain indices for chain-aware geometric constraints.

### Model Architecture

The *Kukulu* model architecture, shown in Figure 1, supports the input of a PDB structure of an antibody-antigen complex. This structure is processed as mentioned above and converted to tensor representations of the prepared and masked structures. Then, the input tensors are fed into their respective encoders. The full model (`StructuralJEPAEngine`) has three trainable components:

**Fig. 1.**
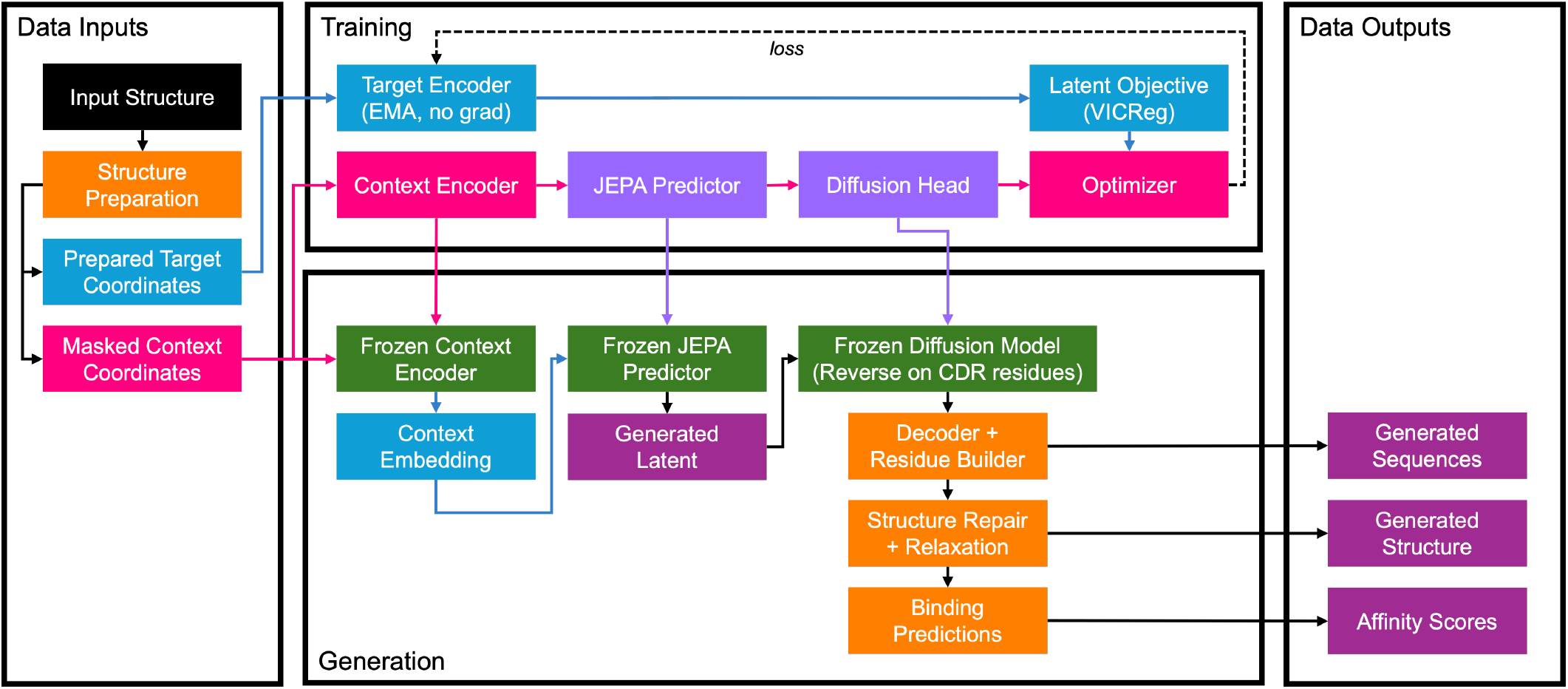
*Kukulu* model architecture showing training and generation pathways.

1. Context/target residue encoder (MLP tokenization of local atom coordinates): coordinates are translated into residue-local frames centered at *C*_*α*_ and embedded into latent tokens.
2. JEPA predictor (transformer encoder): predicts target latents from masked context latents with positional encoding and padding-aware attention.
3. Conditional diffusion head: predicts coordinate noise, atom-presence logits, and residue identity logits for CDR reconstruction.

The target encoder is an exponential moving average (EMA) copy of the context encoder, updated after each optimization step. A backbone-first variant (`BackboneLoopEngine`) uses only N-CA-C-O atoms during diffusion and then reconstructs sidechains from local templates.

### Training Objective

The total loss combines latent prediction, diffusion denoising, and structural reconstruction constraints:

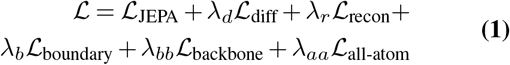

JEPA loss is computed with a masked VICReg objective on CDR tokens only (9), including invariance, variance, and covariance terms. Diffusion uses a Gaussian schedule over timesteps with MSE denoising loss on masked CDR atoms (5).

Reconstruction adds:

1. Atom-presence binary cross-entropy (BCE) loss (`diff`).
2. Coordinate smooth-*L*_1_ loss on supervised atoms (`coord`).
3. Residue identity cross-entropy (`residue`).
4. CDR boundary geometry loss over framework-loop peptide transitions (`boundary`).
5. Backbone bond/angle/dihedral regularization on CDR-local neighborhoods (`backbone`).
6. Intra-residue all-atom pair-distance geometry loss for masked tokens (`atom`).

### Inference and Post-processing

Given an input complex, inference repeats the same preparation and masking logic in memory, then runs reverse diffusion only on masked CDR positions while keeping framework residues fixed. Predicted residues are decoded by argmax or temperature-controlled sampling. Atom masks are thresholded and constrained by residue-specific allowed atom sets.

Optional sidechain rebuilding uses local N-CA-C coordinate frames: per-residue sidechain templates are estimated from framework residues of the same amino acid type, with optional fallback to global templates fit from the training corpus.

Generated structures are exported back to PDB, re-processed with the preparation logic, optionally relaxed with *AMBER*/*OpenMM* routines (14), and scored with *HADDOCK3* for downstream ranking of affinity metrics (17, 18).

### Implementation and Usage Notes

The codebase is implemented in PyTorch and uses modular engines for all-atom and backbone-first generation. Data preparation utilities are in `scripts/prepare/`, tensor conversion in `scripts/convert/`, and inference in `scripts/infer/`. The design API exposes a single `generate()` wrapper, as shown below, for checkpoint-conditioned loop generation from user-provided complexes. This function also performs the PDB repair process (for any missing atoms or residues, and removing unneeded chains) and the side chain relaxation process, if necessary.

~~~
from kukulu import generate
result = generate(
 checkpoint_path=“checkpoint.pt”,
 pdb_path=“1ABC.pdb”,
 h_chain=“H”,
 l_chain=“L”,
 antigen_chains=“A|B”,
 device=“cuda”
)
~~~

This then outputs the generated H and L chain Fv sequences, CDR loop positions, a relaxed PDB string, and predicted binding affinity metrics. An example output for PDB 5jxe is as follows:

~~~
{
 “pdb_id”: “5jxe”,
 “h_chain”: “C”,
 “l_chain”: “D”,
 “antigen_chains”: [“B”],
 “errors”: [],
 …
 “input_h_seq”: “QVQLV…”,
 “input_l_seq”: “EIVLT…”,
 “masked_h_seq”: “…ASXXXXXXXYM…”,
 “masked_l_seq”: “…SCXXXXXXXWY…”,
 “generated_h_seq”: “EIVLT…”,
 “generated_l_seq”: “QVQLV…”,
 “generated_cdr_residue_count”: 55,
 “generated_pdb”: “ATOM …”,
 “relaxation_applied”: true,
 “haddock_scores”: {
  “score”: -217.7198,
  “vdw”: -149.3,
  “elec”: -193.378,
  “desolv”: -29.7442,
  “bsa”: 3823.62,
  “total”: -342.678
 }
}
~~~

### Evaluation

Ten antibody-antigen complexes were selected for benchmarking, shown in Table 1. For each antibody, *Kukulu* was run 500 times, with full relaxation until convergence, to generate unique Fv sequences. Using the resulting affinity scores, the generated structures were compared against the affinity metrics of the respective reference complex. (See Table S1 for reference affinity scores.)

**Table 1.**
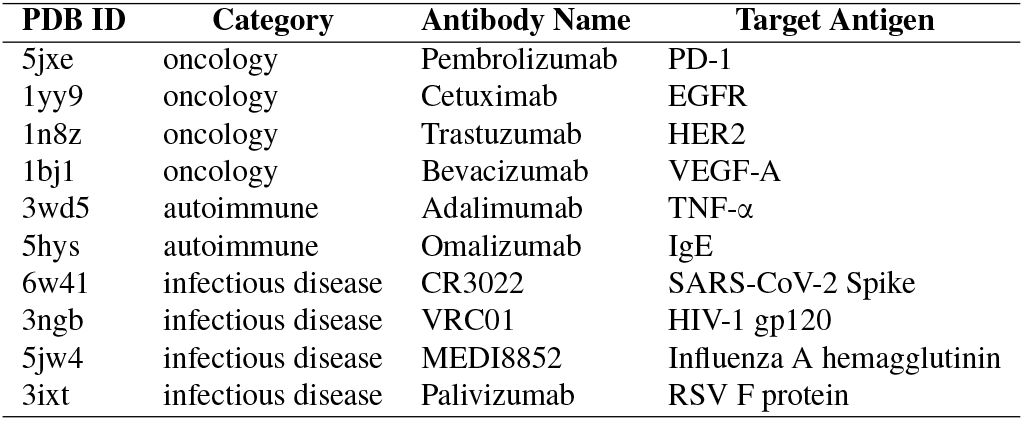
Evaluation antibody-antigen complexes for benchmarking.

| PDB ID | Category | Antibody Name | Target Antigen |
| --- | --- | --- | --- |
| 5jxe | oncology | Pembrolizumab | PD-1 |
| 1yy9 | oncology | Cetuximab | EGFR |
| 1n8z | oncology | Trastuzumab | HER2 |
| 1bj1 | oncology | Bevacizumab | VEGF-A |
| 3wd5 | autoimmune | Adalimumab | TNF-α |
| 5hys | autoimmune | Omalizumab | IgE |
| 6w41 | infectious disease | CR3022 | SARS-CoV-2 Spike |
| 3ngb | infectious disease | VRC01 | HIV-1 gp120 |
| 5jw4 | infectious disease | MEDI8852 | Influenza A hemagglutinin |
| 3ixt | infectious disease | Palivizumab | RSV F protein |

## Results

The *Kukulu* model was trained over 50 epochs on 7,943 samples (paired “prepped” and “masked” structures; 7,148 for training and 795 for validation). Each epoch took around 80 minutes to complete. This achieved an overall training loss of 39.0115 (comprised of the composite losses: jepa=36.0032, diff=0.5902, boundary=0.0925, backbone=0.1668, atom=0.0000, coord=0.0952, and residue=1.9710). Similarly, the overall validation loss was 38.3317 (comprised of: jepa=35.4381, diff=0.5630, boundary=0.0964, backbone=0.1406, atom=0.0000, coord=0.0987, and residue=1.8984).

Across the benchmarking complexes, the 5,000 generated Fv sequences (500 × 10 antibodies) took approximately 26 hours to inference on an NVIDIA RTX 5090.

Overall, *Kukulu* was able to generate many excellent binder candidates, with the best overall scores as follows:

- HADDOCK Score: -504.84
- Total Energy: -1398.89 kcal/mol
- Van der Waals Energy: -258.08 kcal/mol
- Electrostatic Energy: -1222.71 kcal/mol
- Desolvation Energy: -85.39 kcal/mol
- Buried Surface Area: 7496.44 *Å*^2^

Shown in Figure 3, the *Kukulu* model consistently generates novel CDR loops that have desired electrostatic characteristics for the interactions with the epitope. However, enhancements should be made to improve the Van der Waals energies of the interactions as the generated structures often failed to have sufficiently attractive (negative) affinity of this type.

Furthermore, as shown in Table 2, for some benchmark structures such as 5jw4 and 3ngb, there were significantly strong binders generated overall. This includes favorable buried surface area and desolvation energy metrics. In contrast, generated binders for structures such as 1n8z and 1yy9 often failed to beat the reference structures’ binding affinity values. Some examples of improved binding candidates are shown in Figure 2.

**Table 2.** PDB-level metrics of the generated complexes. Values shown are mean values ± standard deviation.

| PDB ID | HADDOCK Score | Van der Waals Energy | Electrostatic Energy | Desolvation Energy | Buried Surface Area | Total Energy |
| --- | --- | --- | --- | --- | --- | --- |
| 5jw4 | -374.46 ± 222.06 | -146.18 ± 216.60 | -903.63 ± 123.94 | -47.55 ± 11.00 | 6560.62 ± 227.26 | -1049.81 ± 274.83 |
| 3ngb | -111.78 ± 234.62 | -9.10 ± 234.16 | -375.02 ± 63.38 | -27.67 ± 9.70 | 3126.46 ± 154.20 | -384.12 ± 250.68 |
| 6w41 | -70.41 ± 236.72 | 14.00 ± 235.58 | -229.82 ± 96.55 | -38.45 ± 11.54 | 2352.02 ± 191.85 | -215.82 ± 259.00 |
| 3wd5 | -32.18 ± 276.22 | 37.85 ± 272.75 | -254.07 ± 100.56 | -19.21 ± 9.87 | 2217.12 ± 172.20 | -216.22 ± 309.32 |
| 3ixt | -17.30 ± 278.61 | 42.67 ± 276.50 | -137.25 ± 77.54 | -32.52 ± 10.63 | 1966.49 ± 244.63 | -94.58 ± 296.52 |
| 5hys | -4.93 ± 284.02 | 58.33 ± 280.64 | -184.28 ± 100.70 | -26.40 ± 11.46 | 2071.95 ± 226.84 | -125.96 ± 316.66 |
| 1bj1 | 21.93 ± 404.05 | 89.55 ± 404.53 | -162.87 ± 76.58 | -35.05 ± 10.25 | 2000.68 ± 160.06 | -73.32 ± 414.78 |
| 5jxe | 197.65 ± 641.13 | 254.95 ± 639.77 | -112.22 ± 70.56 | -34.85 ± 10.24 | 2038.12 ± 184.55 | 142.73 ± 652.01 |
| 1yy9 | 328.18 ± 412.63 | 358.42 ± 414.39 | -80.90 ± 34.49 | -14.07 ± 8.16 | 1504.96 ± 159.77 | 277.53 ± 416.87 |
| 1n8z | 358.91 ± 404.93 | 434.13 ± 404.75 | -222.34 ± 55.02 | -30.76 ± 8.38 | 1805.62 ± 144.14 | 211.80 ± 413.52 |

**Fig. 2.**
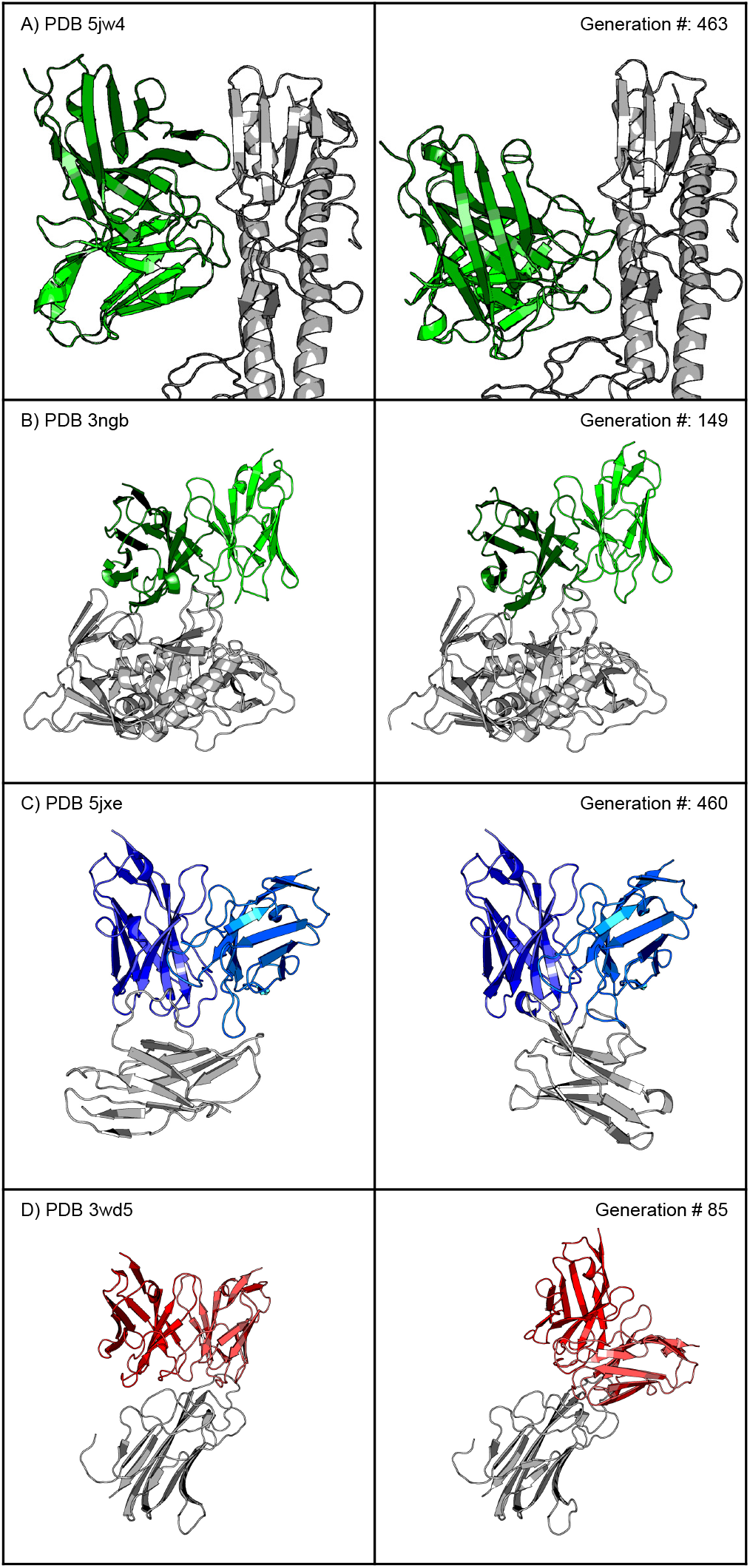
Example comparisons of reference benchmark structures vs. improved binders generated with *Kukulu*.

**Fig. 3.**
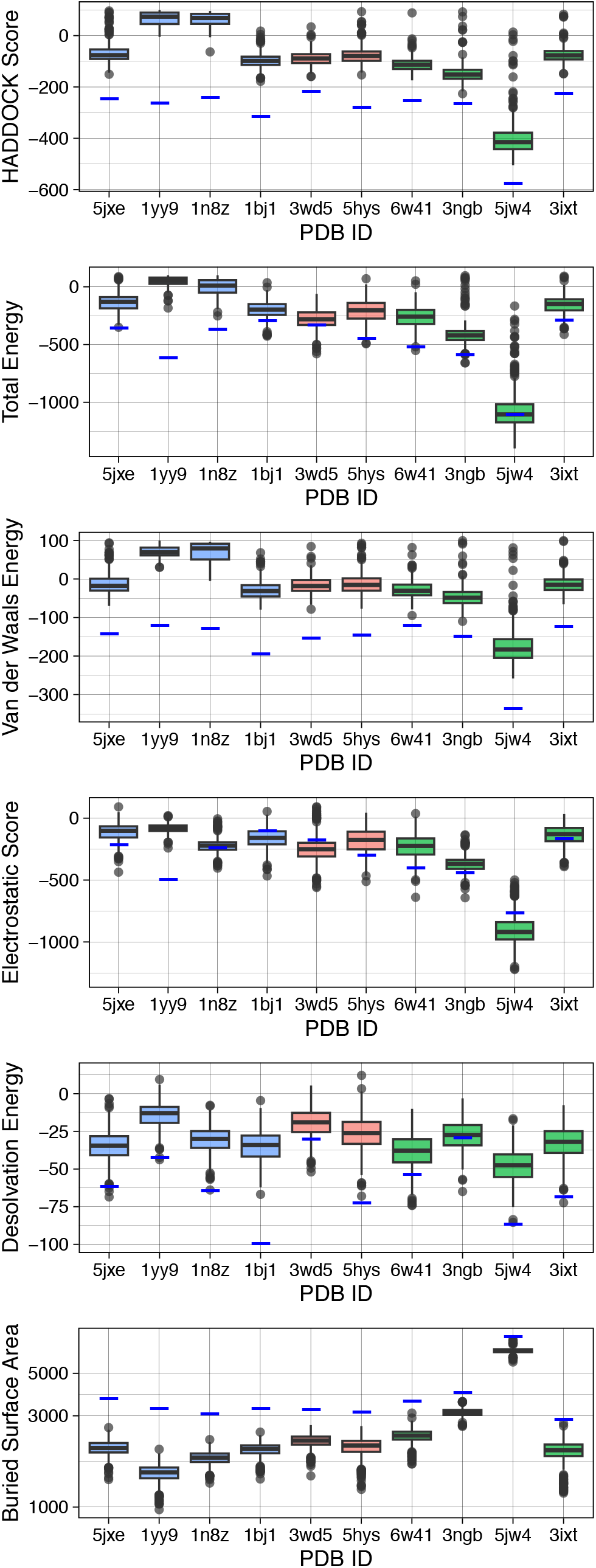
Comparison boxplots of the generated structures vs. reference antibody-antigen complexes (reference metrics shown as blue lines).

## Discussion

*Kukulu* is designed as a practical middle ground between purely energy-based antibody design and unconstrained sequence generation. Relative to classical generative frameworks such as *RosettaAntibodyDesign* (3), *BoltzGen* (19), or *peleke-1* antibody language models (11), this method directly learns CDR structural variability from paired masked/prepared examples.

Relative to global generative diffusion systems (6), *Kukulu* focuses on local loop reconstruction conditioned on fixed framework and antigen context, which is advantageous for iterative lead optimization workflows.

This model is meant to used as part of an existing AI-based antibody discovery workflow to achieve higher variability in the CDR sequence outputs, which may uncover unique therapeutic candidates with desired biophysical characteristics or improved binding over the original structure.

The JEPA + diffusion combination is motivated by representation quality and sampling flexibility. JEPA’s latent prediction encourages context-aware CDR embeddings (8), while diffusion provides stochastic multimodal geometry generation (5). In practice, we found that additional geometry losses are important to avoid chain discontinuities and unrealistic local torsion artifacts at framework-loop boundaries.

Previously, there have been other self-supervised learning models trained using JEPA-based frameworks such as I-JEPA for images (8), V-JEPA for and motion dynamics in video (20), and T-JEPA for tabular data (21). However, our application of JEPA and diffusion is molecular design is novel, both in the realm of embedded architectures in AI-based structural biology more broadly.

Limitations of the present model and approach include sporadic unnatural reconstruction of the CDR loop structures, hence the need for the AMBER relaxation step to move the loops into a lower energy conformation. At present, protein folding may need to be employed to refold the novel Fv sequences in complex with the antigen chains for more accurate scoring. While the CDR sequence reconstruction is reliable, the overall handling of the structure stands to be improved.

Thus, future work should include additional training time with a larger set of input structures and modifications to the structure generation constraints for better output performance. As always with computational modeling, antibody candidates generated with this model need to be validated empirically in future wet lab studies. Future iterations will enable wet-lab-informed evaluation and enhanced postprocessing of the candidates to alert explicit developability constraints, biophysical risks, and manufacturability problems. In future iterations, additional work will be performed to better understand the model’s composite weighted loss and its effect on structure and sequence generation. Also, loss weights must be better calibrated to address the loss imbalance of the JEPA portion that overshadows the structural terms. An ablation matrix is planned to isolate contributions from:

1. JEPA latent objective (with/without VICReg terms).
2. Geometry regularizers (boundary, backbone, all-atom distance losses).
3. Sidechain rebuilding strategy (none, local template, local+global template).
4. Backbone-only versus full atom37 diffusion engines.

These analyses will quantify trade-offs between geometric realism, sequence diversity, and docking-oriented score improvements.

Today, however, *Kukulu* provides as a unique framework for generating novel CDR loop sequences using a structure-aware pipeline with a blended JEPA and diffusion reconstruction approach. With the goal of increasing the creativity and novelty of AI-generated antibodies, *Kukulu* provides a direct generative step that can be easily integrated into existing antibody discovery workflows.

## Acknowledgments

We acknowledge the following entities at the University of North Carolina at Charlotte: the Center for Computational Intelligence to Predict Health and Environmental Risks (CI-PHER), the Department of Bioinformatics and Genomics, and the School of Data Science.

## Code and Data Availability

All code, data, results, and additional analyses are openly available on GitHub at: https://github.com/silicobio/kukulu. This repository includes the open-source logic for data preparation and evaluation for training additional *Kukulu*-like models. Model weights and prepared/-masked PDB structures are hosted on Hugging Face at https://hf.co/collections/silicobio/kukulu.

## Supplementary Materials

**Supplementary Table S1.**
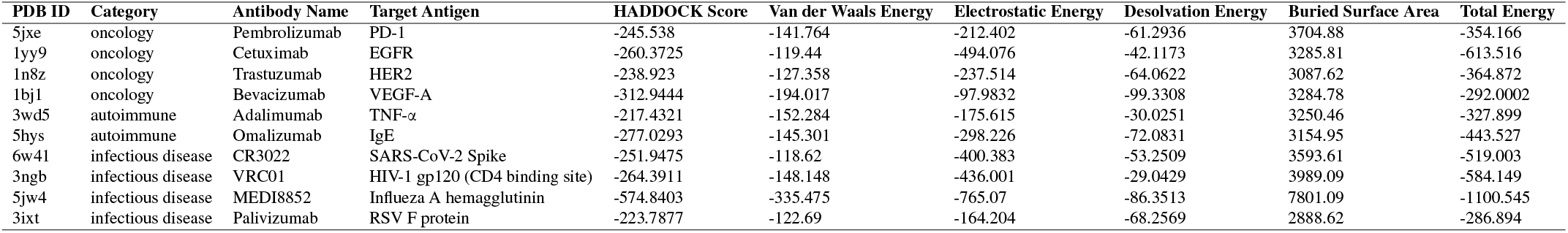
Reference scores of the benchmark antibody-antigen complexes.

| PDB ID | Category | Antibody Name | Target Antigen | HADDOCK Score | Van der Waals Energy | Electrostatic Energy | Desolvation Energy | Buried Surface Area | Total Energy |
| --- | --- | --- | --- | --- | --- | --- | --- | --- | --- |
| 5jxe | oncology | Pembrolizumab | PD-1 | -245.538 | -141.764 | -212.402 | -61.2936 | 3704.88 | -354.166 |
| 1yy9 | oncology | Cetuximab | EGFR | -260.3725 | -119.44 | -494.076 | -42.1173 | 3285.81 | -613.516 |
| 1n8z | oncology | Trastuzumab | HER2 | -238.923 | -127.358 | -237.514 | -64.0622 | 3087.62 | -364.872 |
| 1bj1 | oncology | Bevacizumab | VEGF-A | -312.9444 | -194.017 | -97.9832 | -99.3308 | 3284.78 | -292.0002 |
| 3wd5 | autoimmune | Adalimumab | TNF-α | -217.4321 | -152.284 | -175.615 | -30.0251 | 3250.46 | -327.899 |
| 5hys | autoimmune | Omalizumab | IgE | -277.0293 | -145.301 | -298.226 | -72.0831 | 3154.95 | -443.527 |
| 6w41 | infectious disease | CR3022 | SARS-CoV-2 Spike | -251.9475 | -118.62 | -400.383 | -53.2509 | 3593.61 | -519.003 |
| 3ngb | infectious disease | VRC01 | HIV-1 gp120 (CD4 binding site) | -264.3911 | -148.148 | -436.001 | -29.0429 | 3989.09 | -584.149 |
| 5jw4 | infectious disease | MEDI8852 | Influeza A hemagglutinin | -574.8403 | -335.475 | -765.07 | -86.3513 | 7801.09 | -1100.545 |
| 3ixt | infectious disease | Palivizumab | RSV F protein | -223.7877 | -122.69 | -164.204 | -68.2569 | 2888.62 | -286.894 |

